# Adaptation and response of verrucomicrobial methanotrophs to heat and acidity

**DOI:** 10.1101/2025.09.03.673897

**Authors:** Rob A. Schmitz, Stijn H. Peeters, Theo A. van Alen, Anchelique Mets, Carmen Hogendoorn, Guylaine H.L. Nuijten, Carmen A. Iosif, Arjan Pol, Stefan Schouten, Mike S.M. Jetten, Huub J.M. Op den Camp

## Abstract

Acidophilic microorganisms thrive in environments where the external pH is orders of magnitude lower than their intracellular pH. Verrucomicrobial methanotrophs of the family *Methylacidiphilaceae*, including *Methylacidiphilum* and *Methylacidimicrobium*, inhabit extremely acidic geothermal environments and can grow at a pH < 1.0 and temperatures up to 65 °C. We analyzed and compared their membrane fatty acid compositions at pH 3.0 across strains with different temperature optima. Thermophilic *Methylacidiphilum* strains almost exclusively contain saturated fatty acids, while mesophilic *Methylacidimicrobium* strains incorporate 16–47% unsaturated fatty acids. Notably, the thermophile *Methylacidiphilum fumariolicum* SolV increases unsaturated fatty acid content in response to a 10 °C temperature decrease. Genomic analysis revealed a conserved fatty acid biosynthesis pathway. Despite constitutive expression of predicted pH homeostasis genes, SolV did not upregulate them upon changing the pH from 3.0 to 1.7. However, genes involved in methane oxidation were strongly upregulated, suggesting a potential metabolic adaptation to extreme acidity.

## Introduction

Acidophilic microorganisms thrive in environments that are either naturally acidic (*e*.*g*., acidic geothermal environments and sulfide-rich caves) or acidic due to anthropogenic activity (*e*.*g*., acid mine drainage systems) (Schoen and Rye 1970; Baker and Banfield 2003; Jones et al. 2012). Moderate acidophiles have a pH optimum between 3.0 and 5.0, whereas extreme acidophiles grow optimally below pH 3.0 (Johnson 2007). Prokaryotic acidophiles are found in a range of archaeal and bacterial phyla (Johnson and Quatrini 2020). Methanotrophs of the bacterial phylum Verrucomicrobia (order *Methylacidiphilales*) are found in acidic geothermal environments and grow optimally at a pH of 1.0 to 3.0 (Schmitz et al. 2021a). These harsh environments are located across the globe and are characterized by geothermal features such as mud pools, fumaroles and hot springs from which acidic fluids and gases such as methane (CH_4_), hydrogen gas (H_2_), carbon dioxide (CO_2_) and hydrogen sulfide (H_2_S) are expelled (Castaldi and Tedesco 2005; Picone et al. 2020; Schmitz et al. 2021a; Benson et al. 2011). Such habitats can become extremely acidic if the buffering capacity of the environment is exceeded through microbial oxidation of H_2_S to sulfuric acid (H_2_SO_4_) (Johnson and Quatrini 2020). Interestingly, acidophiles have developed various mechanisms to maintain their intracellular proton concentration several orders of magnitude lower than their surroundings, which is critical to survive in extremely acidic environments (Baker-Austin and Dopson 2007).

Prior to the discovery of verrucomicrobial methanotrophs from acidic geothermal soils, aerobic methanotrophs were thought to be restricted to the Alpha- and Gammaproteobacteria, of which no extreme acidophiles (growth at pH < 3.0) have been isolated (Schmitz et al. 2021a; Whittenbury et al. 1970; Hanson and Hanson 1996). Hereafter, a member of the N10 phylum (*Candidatus* Methylomirabilis oxyfera) was shown to perform aerobic methanotrophy in anoxic environments through an intra-aerobic pathway (Ettwig et al. 2010). Recently, a newly isolated *Mycobacterium* strain from the phylum Actinomycetota was found to oxidize methane at a pH as low as 0.75, expanding the known phylogenetic diversity of aerobic methanotrophs (Kambara et al. 2025).

Currently, it is well established that members of the verrucomicrobial genus *Methylacidiphilum*, which exhibit optimal growth at 50−60 °C and grow up to 65 °C, inhabit acidic geothermal environments across the globe (Dunfield et al. 2007; Pol et al. 2007; Islam et al. 2008; Erikstad et al. 2019; Awala et al. 2021). More recently, verrucomicrobial methanotrophs were isolated from comparable, but significantly cooler environments (Sharp et al. 2014; Van Teeseling et al. 2014). These mesophilic strains, belonging to the genus *Methylacidimicrobium*, exhibit optimal growth at 33−44 °C and grow up to 49 °C, with some able to grow at a pH as low as 0.5 (Van Teeseling et al. 2014). An exception within this mesophilic genus is *Methylacidimicrobium thermophilum* AP8, which grows optimally at 50 °C and up to 55 °C (Picone et al. 2021b). Overall, the primary distinction between *Methylacidiphilum* and *Methylacidimicrobium* appears to be their thermal preference, with the former generally favoring higher temperature.

Interestingly, although all known verrucomicrobial methanotrophs have been isolated from methane-rich acidic geothermal environments, pyrosequencing of samples derived from sulfide-rich concrete sewage pipes has revealed the presence of *Methylacidimicrobium* strains in this non-geothermal habitat (Pagaling et al. 2014). Indeed, recently the verrucomicrobial methanotroph *Methylacidiphilum fumariolicum* SolV was shown to degrade methanethiol (CH_3_SH) and hydrogen sulfide (H_2_S) (Schmitz et al. 2022, 2023). The presence of verrucomicrobial methanotrophs in these man-made environments indicates that they are not limited to acidic geothermal habitats but can also thrive in other acidic niches.

pH homeostasis is essential for all living organisms since a lowered intracellular pH could result in protein instability, reduced enzyme activity, and structural changes of both DNA and RNA (Madshus 1988; Slonczewski et al. 2009). Hence, acidophiles need to maintain a proton gradient of several orders of magnitude between the interior and the external environment (Slonczewski et al. 2009; Moll and Schäfer 1988). Indeed, the verrucomicrobial methanotroph *Methylacidiphilum* sp. RTK17.1 was shown to maintain an intracellular pH of 6.5 when growing in medium with an external pH of 1.5 (Carere et al. 2021). The most convenient adaptation to acid is to possess a membrane with low proton permeability (Driessen et al. 1996). Maintaining such a membrane is especially challenging in thermophiles since with increasing temperature, the membrane becomes increasingly permeable to protons (Driessen et al. 1996). Through homeoviscous adaptation to fluctuations in the environment, microbial membranes can retain the required fluidity for integrity, nutrient permeability and membrane protein functioning (Sinensky 1974). Accordingly, saturated fatty acids decrease membrane fluidity whereas unsaturated fatty acids and branched-chain fatty acids increase membrane fluidity (Denich et al. 2003; Zhang and Rock 2008). The fatty acid chains of thermophiles are generally longer, more saturated and include more branched-chain *iso*-fatty acids, whereas acidophiles possess various adaptations such as a relatively high percentage of branched-chain fatty acids and cyclopropane fatty acids (Siliakus et al. 2017). Previously published fatty acid compositions of nine thermophilic *Methylacidiphilum* strains grown at 55 °C and pH 3.0−3.5 revealed an exceptionally large proportion (on average 99%) of saturated fatty acids (Erikstad et al. 2019; Op den Camp et al. 2009). The high share of saturated fatty acids in *Methylacidiphilum* strains could be an adaptation to establish optimal membrane fluidity even under the combination of a high temperature and an extremely low pH (Konings et al. 2002; Sohlenkamp 2017). Besides having a membrane with low proton permeability, acidophiles seem to share several mechanisms with neutrophilic microorganisms to cope with an excess of protons intracellularly, such as active proton efflux and cytoplasmic buffering (Baker-Austin and Dopson 2007; Foster 2004). While neutrophiles use these mechanisms to swiftly respond to acid stress, acidophiles may use them to live and grow perpetually in acidic environments (Foster 2004; Kanjee and Houry 2013; Guan and Liu 2020).

In this study, we analyzed 11 genomes of different verrucomicrobial methanotrophs to predict how they synthesize membrane fatty acids. Hereafter, we analyzed the fatty acid compositions of three mesophilic verrucomicrobial methanotrophs and compared them with those found in the thermophilic genus *Methylacidiphilum*. In addition, we grew the thermophilic strain *Methylacidiphilum fumariolicum* SolV at different temperatures and pH values in continuous cultures (chemostats) to assess how temperature affects the fatty acid composition and how pH affects gene expression.

## Materials and methods

### Cultivation of *Methylacidiphilum* and *Methylacidimicrobium* strains

*Methylacidiphilum fumariolicum* SolV was grown as methane-limited continuous culture as described before (Schmitz et al. 2020), except that the cells were grown at a pH value of 1.7 and 3.0, at a temperature of 45 and 55 °C and that a 400 mL bioreactor was used. To assess the effect of a temperature and pH decrease, the temperature and pH were decreased abruptly, without the use of a gradient. RNA and fatty acids were extracted and isolated after new steady states were reached after at least three generations. *Methylacidimicrobium tartarophylax* 4AC was grown in a methane-limited chemostat at pH 3.0 and 38 °C as described by Mohammadi et al. (2019). *Methylacidimicrobium cyclopophantes* 3B was grown as batch culture at pH 3.0 and 44 °C and *Methylacidimicrobium fagopyrum* 3C as batch culture at pH 3.0 and 35 °C as described by Van Teeseling et al. (2014). These temperatures were chosen as they are the temperature optima of the different strains (Van Teeseling et al. 2014). To isolate fatty acids, 10 mL biomass was centrifuged at 5000 × *g* at 4 °C for 5 min. The supernatant was decanted and the pellet was resuspended in the remaining liquid by gently pipetting up and down. The cell suspension was transferred to a glass bowl, which was immediately frozen in liquid nitrogen. Finally, the frozen pellet was freeze-dried overnight.

### Fatty acid extraction and analysis

Freeze-dried bacterial biomass was extracted with a modified Bligh and Dyer extraction. The samples were ultrasonically extracted for 10 min with a solvent mixture containing methanol, dichloromethane (DCM) and phosphate buffer (2:1:0.8, vol/vol/vol). After centrifugation, the solvent was collected, combined and the residues were re-extracted twice. A biphasic separation was achieved by adding additional DCM and phosphate buffer to a ratio methanol, DCM and phosphate buffer of 1:1:0.9 (vol/vol/vol). The aqueous layer was washed two more times with DCM and the combined organic layers were dried over a Na_2_SO_4_ column followed by drying under N_2_. The extracts were subsequently hydrolyzed with 1 N 5% HCl in methanol by reflux for 3 h. The hydrolysate was adjusted to pH 4 with 2 N KOH-methanol (1:1, vol/vol) and, after addition of water to a final water-methanol ratio of 1:1, extracted three times with DCM. The DCM fractions were collected and dried over Na_2_SO_4_. The obtained extract was methylated with diazomethane and silylated with *N,O*-bis(trimethylsilyl)trifluoroacetamide (BSTFA). The derivatized extract was analyzed by GC and GC-mass spectrometry (MS). GC-MS was performed using a Triple Quad 7000C GC-MS (Agilent Technologies, Santa Clara, CA USA) in full scan mode. A CP-Sil5 CB column (25 m x 0.32 mm with a film of 0.12 μm, Agilent Technologies) was used for the chromatography with He as carrier gas with a constant flow of 2 mL · min^-1^. The samples (1 μL) were injected onto the column at 70 °C, after which the temperature was increased at 20 °C · min^-1^ to 130 °C, raised further by 4 °C · min^-1^ to 320 °C, at which it was held for 20 min. The fatty acids analyzed in this study of strains *Methylacidimicrobium tartarophylax* 4AC, *Methylacidimicrobium cyclopophantes* 3B, *Methylacidimicrobium fagopyrum* 3C and *Methylacidiphilum fumariolicum* SolV were compared with fatty acids of *Methylacidiphilum kamchatkense* Kam1 and *Methylacidiphilum infernorum* V4 as reported by Op den Camp et al. (2009).

### Genetic analyses

To investigate how membrane fatty acids could be synthesized in verrucomicrobial methanotrophs, the following amino acid sequences of enzymes of *Escherichia coli* strains were retrieved from GenBank (Clark et al. 2016): AccA (CAD6021894.1), AccB (CAD6001842.1), AccC (CAD6001830.1), AccD (CAD6008109.1), AcpP (CAD6016753.1), AcpS (CAD6006565.1), FabD (CAD6010950.1), FabH (CAD6016771.1), IlvB (CAD5998393.1), IlvH (CAD6022235.1), IlvC (CAD6023092.1), IlvD (CAD6021386.1), IlvE (CAD6023096.1), LeuA (CAD6022244.1), LeuB (CAD6022247.1), LeuC (CAD6022250.1), LeuD (CAD6022253.1), PdhA (WP_214096071.1), PdhB, (WP_151040350.1), PdhC (ADX52374.1), Pld (WP_216348498.1), FabG (EGT67884.1), FabZ (CAD6021915.1), FabI (CAD6015223.1) and FabF (CAD6016748.1). Subsequently, these sequences were blasted against the following TaxIds in NCBI (Edgar 2010): Methylacidiphilales (taxid: 717963), *Methylacidiphilum* (taxid: 511745), *Methylacidiphilum fumariolicum* SolV (taxid: 1156937), *Methylacidiphilum kamchatkense* Kam1 (taxid: 431057), *Methylacidiphilum* sp. RTK17.1 (taxid: 1776078), *Methylacidiphilum* sp. Yel (taxid: 1847730), *Methylacidiphilum* sp. Phi (taxid: 1847729), *Methylacidiphilum infernorum* V4 (taxid: 481448), *Methylacidimicrobium tartarophylax* (taxid: 1041768), *Methylacidimicrobium cyclopophantes* (taxid: 1041766), *Methylacidimicrobium fagopyrum* (taxid: 1041767), *Verrucomicrobium* sp. LP2A (taxid: 478741), Verrucomicrobia bacterium 4ac (taxid: 1041768), *Verrucomicrobium* sp. 3C (taxid: 1134055), *Methylacidimicrobium thermophilum* A8 (taxid: LR797830) and *Methylacidithermus pantelleriae* PQ17 (taxid: GCA_905250085). BLOSUM62 was used as matrix to compare sequences with an e-value of 10^-6^.

### Core gene cluster analysis

Protein coding genes from assemblies of verrucomicrobial methanotrophs where clustered using usearch (Edgar 2010), with a 0.5 cutoff. These clusters were visualized using the pheatmap package (Kolde 2015) in the R environment (version 4.5.1, R Core Team, 2025).

### RNA isolation, RNA-seq and data analysis

*M. fumariolicum* SolV cells (OD_600_ ~ 2.0) were grown in a chemostat at pH 3.0 and subsequently at pH 1.7. Per condition, three technical replicates consisting of 10 mL cells were collected from the chemostat to create triplicates for transcriptomics. Cells were immediately pelleted for 3 min at 15,000 × *g*, snap-frozen in liquid nitrogen and stored at −80 °C. Total RNA was isolated using the RiboPure™ RNA Purification Kit for bacteria (Thermo Fisher Scientific, Waltham, MA, USA) according to the manufacturer’s protocol. Ribosomal RNA was removed from the total RNA samples to enrich for mRNA using the MICROBExpress Bacterial mRNA Enrichment Kit (Thermo Fisher Scientific) according to the manufacturer’s protocol. The Qubit high sensitivity RNA and the Agilent RNA 6000 Nano kits and protocols were used for the quantitative and qualitative analysis of the extracted total RNA and enriched mRNA. The latter was used for library preparation by using the TruSeq Stranded mRNA Reference Guide (Illumina, San Diego, CA, USA) according to the manufacturer’s protocol. For quantitative and qualitative assessment of the synthesized cDNA, the Qubit double stranded DNA High Sensitivity and the Agilent High Sensitivity DNA kits and protocols were used.

Transcriptome reads were checked for quality using FastQC (Andrews 2010) and subsequently trimmed 10 base pairs at the 5’ end and 5 base pairs at the 3’ end of each read. Reads were mapped against the strain SolV Mfumv_2 genome (accession number LM997411) using Bowtie2 (Langmead and Salzberg 2012). The remainder of the analysis and the production of images were done in the R environment (version 4.5.1, R Core Team, 2025). The mapped read counts per gene were determined using Rsubread (Liao et al. 2019) and fold change and dispersion were estimated using DEseq2 (Love et al. 2014). Before doing any statistics, a principal component analysis on the top 1000 genes by variance of each sample was performed to check whether samples within the same condition were both similar to each sample part of the same condition and dissimilar to any other sample. For differential expression, a Wald test was employed by DEseq2 to calculate adjusted p-values. Differences in counts were considered to be significant if the base mean was higher than 4, the log2 fold change was higher than [0.58], and the adjusted p-value was ≤ 0.05. For easy comparisons between samples, we calculated TPM (Transcripts Per Million) values.

## Results and Discussion

### Verrucomicrobial methanotrophs possess a conserved fatty acid biosynthesis pathway

Based on genomic analyses and comparisons with fatty acid biosynthesis pathways in diverse microorganisms, the fatty acid biosynthesis pathway in verrucomicrobial methanotrophs could be predicted (**Supplementary File S1**). Although several differences are found between the genera *Methylacidiphilum* and *Methylacidimicrobium*, their fatty acid biosynthesis pathways show strong similarity to the well-described pathway of *Escherichia coli* (Zhang and Rock 2008). The fatty acid biosynthesis in verrucomicrobial methanotrophs is thought to be initiated by the conversion of acetyl-CoA to malonyl-CoA by the acetyl-CoA carboxylase (Cronan and Waldrop 2002). The enzyme involved is encoded by the genes *accABCD*, which are not organized in an operon (**Supplementary File S1**). Subsequently, the malonyl-group of malonyl-CoA could be transferred to the acyl carrier protein (ACP) by the enzyme FabD to form malonyl-ACP (Ruch and Vagelos 1973). ACP is typically activated by the ACP synthase and subsequently all intermediates in the fatty acid biosynthesis pathway are attached to this carrier protein (Marcella et al. 2017). The enzyme FabH could condense malonyl-ACP with an acyl-CoA molecule, leading to the formation of a β-ketoacyl-ACP molecule (Jackowski and Rock 1987). FabH has been shown to utilize acyl-CoA primers of different lengths, which results in the synthesis of fatty acids of different acyl chain lengths (Khandekar et al. 2001; Qiu et al. 2005). After the synthesis of β-ketoacyl-ACP, the elongation phase of the fatty acid biosynthesis pathway is initiated, in which verrucomicrobial methanotrophs are predicted to use well-studied enzymes. FabG is known to oxidize an NADPH molecule and reduce the product of FabH to β-hydroxy-acyl-ACP (Lai and Cronan 2004). Subsequently, this product is hydrolyzed to form *trans*-2-enoyl-ACP. In verrucomicrobial methanotrophs, a predicted bifunctional enzyme LpxC/FabZ is encoded that might catalyze this reaction. The final step in the elongation cycle to produce acyl-ACP is presumably catalyzed by the enoyl-ACP reductase FabI in verrucomicrobial methanotrophs. In other microorganisms, the isoforms FabV, FabK and FabL with a similar function have been found as well (Rana et al. 2020). In order to add two additional C-atoms to the acyl-chain, the cycle could rerun, condensing another malonyl-group of malonyl-CoA with acyl-ACP. This condensation reaction is presumably catalyzed by FabF in verrucomicrobial methanotrophs (Wang et al. 2004; **Supplementary File S1**).

Saturated fatty acids to which a methyl-group is attached to the acyl chain increase membrane fluidity compared to non-methylated saturated fatty acids (Kaneda 1991). In turn, a methyl-group attached in *anteiso*-fashion increases membrane fluidity compared to *iso*-fatty acids (Zhang and Rock 2008). Indeed, in *Methylacidiphilum* strains various *anteiso*- and *iso*-fatty acids were previously found (Erikstad et al. 2019; Op den Camp et al. 2009), which could be an adaptation to both a low pH and high temperature (Siliakus et al. 2017; Santiago et al. 2012). FabH, the enzyme that initiates the elongation phase, utilizes an acyl-CoA molecule as primer to produce straight-chain fatty acids. Alternatively, FabH can utilize branched-chain acyl-CoA primers to produce *anteiso*- and *iso*-fatty acids (Qiu et al. 2005). These primers are formed through modification of branched-chain amino acids and are subsequently modified through the same elongation cycle as for the synthesis of straight-chain fatty acids (Kaneda 1991; Choi et al. 2000; Li et al. 2005; Singh et al. 2009; Yu et al. 2016). Verrucomicrobial methanotrophs possess the gene *ilvE* encoding a branched-chain aminotransferase (Santiago et al. 2012). Depending on the type of *anteiso*- and *iso*-fatty acid to be synthesized, this enzyme can convert valine to 3-methyl-oxobutyric acid, leucine to 4-methyl-oxopentanoic acid and isoleucine to 3-methyl-oxopentanoic acid, respectively (Qiu et al. 2005). Subsequently, the branched-chain α-keto acid dehydrogenase (BKD complex) can convert 3-methyl-oxobutyric acid to isobutyryl-CoA, 4-methyl-oxopentanoic acid to isovaleryl-CoA and 3-methyl-oxopentanoic acid to 2-methyl-butyryl-CoA, respectively. These molecules are the precursors that could be used by FabH to produce branched-chain fatty acids and subsequently enter the elongation cycle (Qiu et al. 2005). The even-numbered *iso*-C14:0 and *iso*-C16:0 fatty acids are formed with isobutyryl-CoA as primer, whereas odd-numbered *iso*-C15:0 and *iso*-C17:0 fatty acids are formed by using isovaleryl-CoA. In addition, odd-numbered fatty acids *anteiso*-C15:0 and *anteiso*-C17:0 originate from the primer 2-methyl-butyryl-CoA.

### *Methylacidiphilum* and *Methylacidimicrobium* strains possess dissimilar fatty acid profiles

The large percentage of saturated fatty acids of thermophilic *Methylacidiphilum* strains could be an adaptation to the extreme environment in which these methanotrophs live. To assess the effect of temperature on fatty acid composition, three mesophilic *Methylacidimicrobium* strains were grown at their optimal temperatures (35– 44 °C) and pH 3.0, and their fatty acid profiles were analyzed and compared to those of three thermophilic *Methylacidiphilum* strains grown at their respective optima of 55 °C and pH 3.0 (Op den Camp et al. 2009). The most common fatty acids detected in the mesophilic *Methylacidimicrobium* strains (with an average percentage >10%) are, in sequential order, stearic acid (C18:0), oleic acid (C18:1 (Δ9)), *anteiso*-pentadecanoic acid (aC15:0) and palmitic acid (C16:0) (**Table 1**). Strain 3C and strain 4AC possess similar fatty acid profiles whereas strain 3B produces various other fatty acids in high amounts. In strain 3B, the fatty acid aC15:0 is only present in low quantity (3%), while it possesses a relatively high amount of various unsaturated fatty acids (47%) and hydroxy-fatty acids (10%). Analysis of the fatty acids of *Methylacidiphilum fumariolicum* SolV retrieved in this study show some differences to the fatty acids retrieved from this bacterium several years ago (Op den Camp et al. 2009). These differences may be caused by the fact that in this study, *M. fumariolicum* SolV was grown in a chemostat instead of in batch cultures. Furthermore, in this study the fatty acids were released by acid hydrolysis while previously base hydrolysis was used, which may induce differences in fatty acid composition (Lambert and Moss 1983).

**Table 1:** Percentages of identified fatty acids of three mesophilic verrucomicrobial methanotrophs and one thermophilic (strain SolV) verrucomicrobial methanotroph.

| Fatty acid | strain 3B | strain 3C | strain 4AC | strain SolV |
| --- | --- | --- | --- | --- |
| C14:0 | – | 0.2 | 1.9 | 3.4 |
| iso-C14:0 | – | 1.6 | 6.4 | 9.4 |
| C15:0 | – | – | – | 1.7 |
| anteiso-C15:0 | 2.9 | 25.9 | 22.9 | 17.6 |
| iso-C15:0 | – | 0.2 | 0.7 | – |
| C16:0 | 10.1 | 13.8 | 17.9 | 17.5 |
| iso-C16:0 | 1.1 | 2.7 | 2.3 | 5.4 |
| C16:0 (3-OH) | – | – | – | 0.1 |
| C17:0 | 3.9 | 0.5 | 0.3 | 0.5 |
| anteiso-C17:0 | – | 4.9 | 1.2 | 1.4 |
| iso-C17:0 | – | 0.9 | 0.7 | 0.3 |
| C18:0 | 31.4 | 29.7 | 28.9 | 41.8 |
| iso-C18:0 | – | – | – | 0.9 |
| C18:0 (3-OH) | 3.6 | 0.4 | 0.6 | 0.1 |
| C18:1 (unknown position) | – | – | 0.4 | – |
| C18:1 ($\Delta 9$) | 26.3 | 16.4 | 14.8 | – |
| C18:1 ($\Delta 11$) | 8.4 | 1.3 | 1.0 | – |
| C18:1 (6-OH + 8-OH) | 5.9 | 0.7 | – | – |
| C19:1 (unknown position) | – | 0.8 | – | – |
| C19:1 ($\Delta 9$, $\Delta 10$ & $\Delta 13$) | 6.3 | – | – | – |

Strain 3B: *Methyacidimicrobium cyclopophantes* 3B (grown at 44°C and pH 3.0 in batch); strain 3C: *Methyacidimicrobium fagopyrum* 3C (grown at 35°C and pH 3.0 grown in batch); strain 4AC: *Methyacidimicrobium tartarophylax* 4AC (grown at 38°C and pH 3.0 grown in a chemostat); strain SolV: *Methyacidiphilum fumariolicum* SolV (grown at 55°C and pH 3.0 in a chemostat)

Whereas the thermophilic *Methylacidiphilum* strains almost exclusively possess saturated fatty acids, the membranes of the mesophilic *Methylacidimicrobium* strains consist on average for 27% of unsaturated fatty acids (**Fig. 1a**). This difference is not statistically significant due to substantial variation in membrane composition between species within the genus *Methylacidimicrobium* (Welch’s two-sample t-test, p = 0.11). The minimal proportion of unsaturated fatty acids in thermophilic *Methylacidiphilum* strains is consistent with the general trend that microbes growing at higher temperatures contain lower levels of unsaturated fatty acids in comparison with those thriving at lower temperatures, to maintain proper membrane fluidity (Suutari and Laakso 1994). The membranes of *Methylacidiphilum* strains consist for 19% ± 10% of *iso*-fatty acids, compared to 6% ± 5% in the *Methylacidimicrobium* strains (Welch’s two-sample t-test, p = 0.13). A high proportion of *iso*-fatty acids is typical for thermophiles, whereas branched-chain fatty acids in general are typically detected in high amounts in acidophiles (Siliakus et al. 2017; Oshima and Miyagawa 1974). Another general phenomenon in thermophiles is the longer acyl chain of fatty acids compared to microorganisms growing at lower temperatures (Siliakus et al. 2017). The longer the acyl chain the more interaction between neighboring acyl chains of fatty acids and hence an increase in rigidity (Russell 1989). Surprisingly, however, the thermophilic *Methylacidiphilum* strains possess shorter acyl chain lengths compared to their mesophilic counterparts (**Fig. 1b**). Almost 60% (in abundance) of the fatty acids detected in *Methylacidimicrobium* strains have an acyl chain length of at least 18 carbon atoms, whereas shorter fatty acids are more common in the *Methylacidiphilum* strains.

**Fig. 1.**
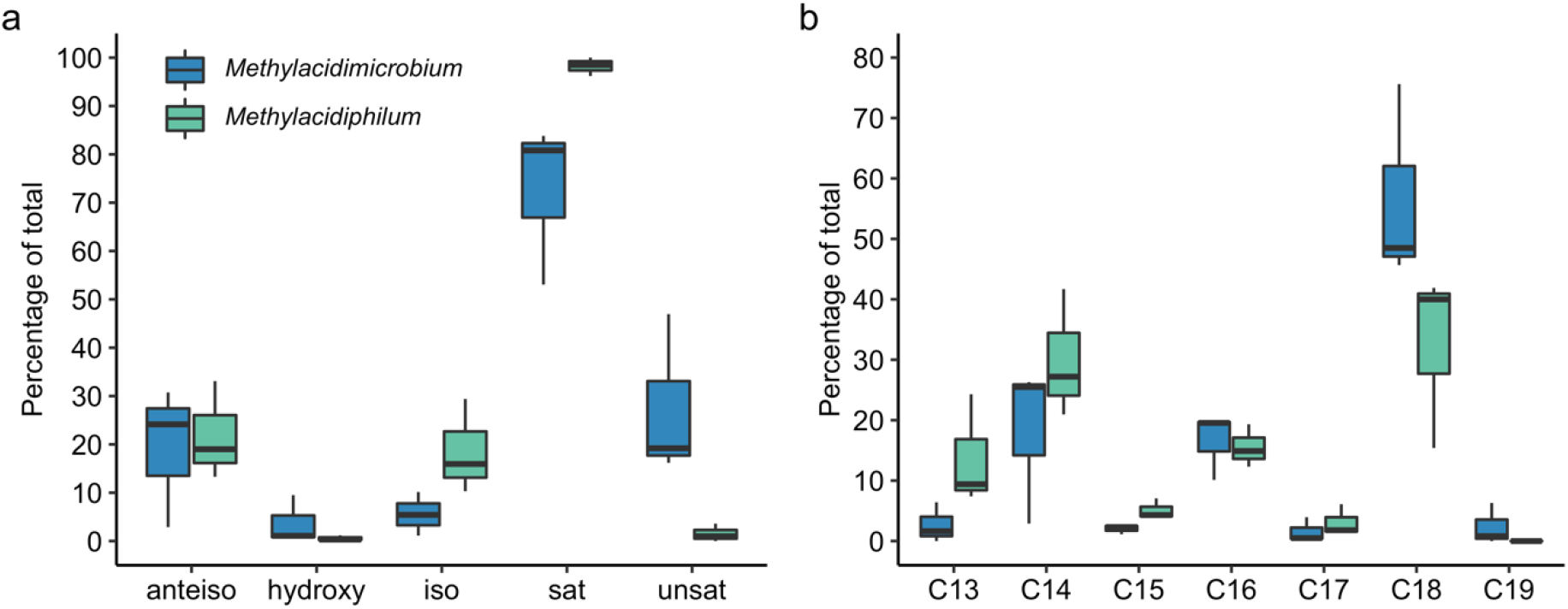
(**a**) Boxplots of different types of fatty acids and (**b**) of different chain lengths detected in three mesophilic *Methylacidimicrobium* strains (*Methylacidimicrobium tartarophylax* 4AC, *Methylacidimicrobium cyclopophantes* 3B *Methylacidimicrobium fagopyrum* 3C) and three *Methylacidiphilum* strains (*Methylacidiphilum fumariolicum* SolV, *Methylacidiphilum kamchatkense* Kam1 and *Methylacidiphilum infernorum* V4). anteiso: anteiso fatty acids; hydroxy: hydroxyl fatty acids; iso: iso-fatty acids; sat: saturated fatty acids; unsat: unsaturated fatty acids. The C-number indicates the length of the acyl chain of the fatty acids, without taking methyl-groups into account

### *M. fumariolicum* SolV synthesizes unsaturated fatty acids at lower temperature

In the natural, environment, verrucomicrobial methanotrophs could encounter rapid fluctuations in temperature and would therefore need to modify their fatty acid composition accordingly to maintain the desired membrane fluidity. To investigate the effect of temperature on fatty acids, *Methylacidiphilum fumariolicum* SolV was grown in a chemostat at 55 °C and subsequently abruptly at 45 °C, after which the fatty acids were analyzed when the culture reached steady state. *M. fumariolicum* SolV reacts to a decrease in growth temperature of 10 °C by synthesizing the unsaturated fatty acid C18:1, which is not detected when grown at 55 °C (**Supplementary Figure 1**). This monounsaturated fatty acid increases the membrane fluidity, which is a common strategy in response to a decrease in temperature (Siliakus et al. 2017). Indeed, the mesophilic *Methylacidimicrobium* strains that were grown at temperatures of 35−44 °C possess large proportions of monounsaturated C18 fatty acids (**Table 1**). However, the membrane fatty acid composition of *M. fumariolicum* SolV grown at pH 3.0 and 45 °C is still distinct from the fatty acid composition of *Methylacidimicrobium cyclopophantes* 3B grown at pH 3.0 and 44 °C. The former strain was grown in a continuous chemostat while the latter was grown in batch at maximum specific growth rate (µ_max_), which may influence the fatty acid composition of the membrane. In general, closely related microorganisms cultivated under the same extreme conditions can have various ways to achieve the necessary membrane fluidity and integrity, which is observed in a variety of acidophiles (Siliakus et al. 2017).

Based solely on genomic comparisons, it cannot be concluded how unsaturated fatty acids are produced by verrucomicrobial methanotrophs *de novo*. Alpha- and Gammaproteobacteria possess a gene encoding FabA, catalyzing the same dehydration reaction in the elongation cycle as FabZ, which these Proteobacteria also possess (Zhang and Rock 2008). Unlike FabZ, FabA is also able to convert *trans*-2-decenoyl-ACP (*i*.*e*., a *trans*-2-enoyl-ACP molecule with 10 carbons) to *cis*-3-enoyl-ACP, creating the precursor for the synthesis of *cis*-unsaturated fatty acids (Heath and Rock 1996; Dodge et al. 2019). FabA therefore represents a branch point between the elongation of saturated fatty acids and the synthesis of unsaturated fatty acids. In *Methylacidimicrobium* strains, but not in *Methylacidiphilum* strains, a gene is found that is classified by InterPro as part of the family “Beta-hydroxydecanoyl thiol ester dehydrase FabA/FabZ” (IPR013114). Accordingly, the *Methylacidimicrobium* strains might use this enzyme for the synthesis of unsaturated fatty acids, which are present only in very low quantity in *Methylacidiphilum* strains (Erikstad et al. 2019; Op den Camp et al. 2009). However, to elongate unsaturated fatty acids produced via FabA, the enzyme FabB is required as well, which is absent in all verrucomicrobial methanotrophs (Heath and Rock 1996; Campbell and Cronan 2001).

Mutation studies in *E. coli* and *Pseudomonas aeruginosa* strains revealed FabA to be essential for the synthesis of unsaturated fatty acids (Wang and Cronan 2004). Interestingly, however, in the Gram-positive bacterium *Enterococcus faecalis*, two FabZ homologs and two FabF homologues are found (Wang and Cronan 2004). Complementation studies revealed that one of the FabZ and FabF homologs of *E. faecalis* can take over the function of FabA and FabB in *E. coli*, respectively. Thus, FabA and FabB are not necessarily required for *de novo* synthesis of unsaturated fatty acids in all bacteria. In this light it is interesting to note that only *Methylacidimicrobium* strains possess two dissimilar homologues of FabZ and FabF, whereas the *Methylacidiphilum* strains only possess one of each. Nevertheless, Wang and Cronan (2004) noted that amino acid sequence comparisons do not suffice in functional predictions, since one FabZ homolog of *E. faecalis* functions as a FabA enzyme but is much more similar in amino acid sequence identity to *E. coli*-FabZ than to *E. coli*-FabA.

The fact that *Methylacidimicrobium* strains possess large amounts of unsaturated fatty acids whereas in *Methylacidiphilum* strains they were barely detected, combined with the additional genes encoding FabZ and FabF homologs in *Methylacidimicrobium* strains that are absent in *Methylacidiphilum* strains, could point to a specific regulation for unsaturated fatty acid synthesis. Nevertheless, mutagenesis studies and isolation of these enzymes are needed for verification. Apart from the *de novo* synthesis of unsaturated fatty acids, microorganisms possess mechanisms to desaturate straight-chain fatty acids by introducing a *cis*-bond in response to environmental changes such as a decrease in temperature (Mansilla et al. 2004; Aquilar and de Mendoza 2006). Interestingly, both *Methylacidiphilum* and *Methylacidimicrobium* strains encode for desaturases to catalyze this reaction, but of different types (**Supplementary File S1**). A temperature decrease from 55 °C to 45 °C in *M. fumariolicum* SolV led to a reduction in C18:0 and a corresponding increase in C18:1 membrane fatty acids, suggesting that a desaturase may have catalyzed this conversion as a mechanism to respond to a sudden temperature decreases (**Supplementary Figure S1**).

Little is known about changes in fatty acid composition as a response to acid stress in acidophiles. The acidophilic sulfur-oxidizer *Acidithiobacillus ferrooxidans* responds to a decrease in pH by an increase in the ratio saturated/unsaturated fatty acids (Mykytczuk et al. 2010). In addition, microorganisms were shown to convert monounsaturated fatty acids to cyclopropane fatty acids in response to a decrease in pH, catalyzed by a cyclopropane fatty acids synthase (Zhang and Rock 2008; Sohlenkamp 2017). However, the gene encoding this enzyme is absent in verrucomicrobial methanotrophs. Acidophiles seem to synthesize various combinations of fatty acids to grow in an environment with low pH, such as *iso*- and *anteiso*-branched chain fatty acids and β-hydroxy fatty acids (Siliakus et al. 2017). Nevertheless, acidophiles can have very distinct fatty acid compositions and therefore multiple mechanisms to thrive at low pH seem to exist.

Gram-negative bacteria such as verrucomicrobial methanotrophs possess a cytoplasmic bilayer composed of various phospholipid fatty acids and an outer membrane composed of phospholipids in the inner leaflet and polysaccharides in the outer leaflet (Speth et al. 2012; Konovalova et al. 2017; Vergalli et al. 2020). Cells of *Escherichia coli* that suddenly encountered a lower pH in the environment were observed to have a similarly low pH in the periplasm, suggesting that the outer membrane is relatively permeable to protons compared to the cytoplasmic membrane (Wilks and Slonczewski 2007). The outer membranes of acidophiles could either be less permeable to protons or they could possess periplasmic proteins that are adapted to a low pH. Interestingly, the purified periplasmic lanthanide-dependent methanol dehydrogenase XoxF of *Methylacidiphilum fumariolicum* SolV has an optimum activity at neutral pH, which could imply a relatively proton-impermeable outer membrane (Keltjens et al. 2014). In addition to this, verrucomicrobial methanotrophs do not possess genes encoding the periplasmic chaperones HdeA and HdeB found in many gram-negative neutrophiles to prevent protein aggregation at low pH (Tapley et al. 2009; Hong et al. 2012). The fatty acid analyses of different thermophilic verrucomicrobial methanotrophs revealed fatty acid profiles with large amounts of saturated fatty acids, which represents a combination of fatty acids in the cytoplasmic membrane and the outer membrane (Erikstad et al. 2019).

### Putative pH homeostasis mechanisms were detected in the genomes

To investigate whether verrucomicrobial methanotrophs possess genes that could be involved in pH homeostasis, a two-pronged approach was used. Firstly, the genomes of five *Methylacidiphilum* strains, five *Methylacidimicrobium* strains and the metagenome-derived genome of *Methylacidithermus pantelleriae* PQ17, representing a third genus of verrucomicrobial methanotrophs, were investigated for shared gene clusters. 407 core gene clusters were found, of which 403 could be assigned a pfam domain using the gathering threshold (**Fig. 2**). These gene clusters were used to search for genes involved in pH homeostasis. In general, large numbers of gene clusters are shared between verrucomicrobial methanotrophs, although clear differences between genera were found (**Fig. 2**). Secondly, genes known or predicted to be involved in pH homeostasis in acidophiles and genes encoding proteins known or predicted to respond to a sudden decrease in pH in neutrophilic microorganisms were BLASTed against the genomes of the verrucomicrobial methanotrophs.

**Fig. 2.**
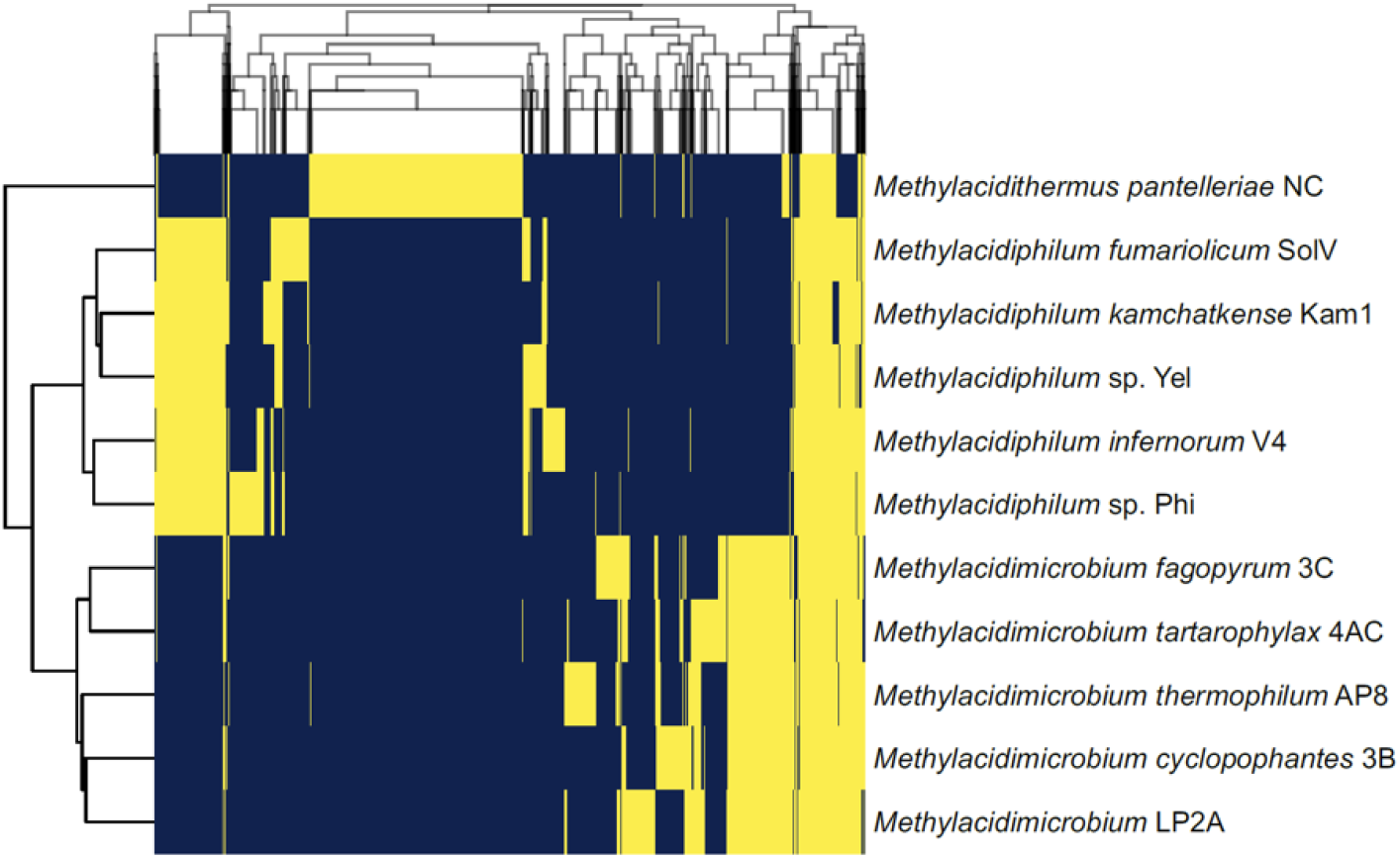
Absence and presence of gene clusters in verrucomicrobial methanotrophs. The core genome consists of 407 gene clusters, out of a total of 7326 detected gene clusters. Genes are assigned to gene clusters using usearch, with a 0.5 cutoff (in yellow > 0.5, in blue < 0.5)

The most convenient mechanism for pH homeostasis is preventing protons from entering the cell, which can be achieved by synthesizing a membrane with relatively low proton permeability (Konings et al. 2002). Still, protons enter the cell via the ATP synthase at the expense of the proton motive force. This force is made up of a membrane potential (ΔΨ) and a pH gradient (ΔpH), creating an energized membrane. In acidophiles, ΔpH is very large due to the acidic environment and the circumneutral interior pH. Under acid stress, *Escherichia coli* was shown to invert the membrane potential to limit proton influx, resulting in a net positive charge close to the inside of the cytoplasmic membrane compared to the outside of the membrane (Baker-Austin and Dopson 2007; Matin et al. 1982; Zychlinsky and Matin 1983; Richard and Foster 2004). Experiments with acidophilic archaea of the genus *Sulfolobus* suggest this inverted electrical gradient is obtained by the influx of potassium (K^+^) ions via dedicated transporters, diminishing the entry of protons over the membrane (Buetti-Dinh et al. 2016). Indeed, *Methylacidiphilum* sp. RTK17.1 was shown to maintain a reversed electrical gradient at a range of pH values (Carere et al. 2021). In verrucomicrobial methanotrophs, the operon *kdpABC* is found (**Supplementary File S2**). This operon is present in a variety of microorganisms and encodes a high-affinity potassium-transporting ATPase involved in potassium uptake (Epstein 2003; Pedersen et al. 2019, Neira et al. 2022). When *E. coli* is deprived of potassium, the sensor kinase KdpD and response regulator KdpE activate the expression of the potassium-translocating ATPase, which are also found in verrucomicrobial methanotrophs (Altendorf et al. 1994; Heermann and Jung 2010). However, in the genome of *M. pantelleriae* PQ17 this potassium-transporting system could not be detected. Since this genome was derived from a metagenome and no isolate is available, it is unknown at what pH range this bacterium lives and therefore whether an inverted electrical gradient is necessary (Picone et al. 2021a). Nonetheless, *M. pantelleriae* PQ17 does seems to encode for a dedicated potassium transport channel, as do the other verrucomicrobial methanotrophs. In addition, several verrucomicrobial methanotrophs encode for a protein annotated as low-affinity kup-type potassium transporter (**Supplementary File S2**).

Nutrients and solutes enter the cell via primary or secondary transporters. Through the latter, acidophiles can make use of the influx of protons along the large pH gradient. Protons that have entered the cell either through transporters, leakage or through the ATP synthase, also need to be removed from the cell to prevent acidification (Baker-Austin and Dopson 2007). Verrucomicrobial methanotrophs possess two types of proton-translocating ATPases used to conserve energy through respiration (Schmitz et al. 2021a; Kruse et al. 2019). Since these protein complexes can work in both directions, they might be used to extrude protons (Slonczewski et al. 2009; Hou et al. 2008). In addition, Kruse et al. (2019) detected two possible sodium/proton antiporters KefB and a sodium/proton-translocating pyrophosphatase HppA that could all be involved in proton extrusion and are also found in other verrucomicrobial methanotrophs, except for *M. pantelleriae* PQ17 (**Supplementary File S2**). Genes encoding enzymes with the same proton-pumping mechanisms were found in the genomes of proteobacterial methanotrophs that live in acidic forest soil at a pH of 3 to 4 (Nguyen et al. 2018).

*Methylacidiphilum* sp. RTK17.1 was shown to maintain an intracellular pH of 6.5 at an extracellular pH of 1.5 to 3.0 (Carere et al. 2021). When its electron transport chain was uncoupled with the use of the chemicals valinomycin and nigericin, the internal pH only decreased to 6.1, which may be the result of the buffering capacity of the cytoplasm and the strong membrane barrier (Carere et al. 2021). In neutrophilic microorganisms, many proteins involved in intracellular proton consumption are upregulated under acid stress (Slonczewski et al. 2009). Verrucomicrobial methanotrophs possess two genes encoding enzymes typically involved in intracellular proton consumption (**Supplementary File S2**). The glutamate decarboxylase (GadB) is a cytoplasmic enzyme found in many microorganisms to cope with acute acid stress (Lin et al. 1995; Lund et al. 2014). The enzyme catalyzes the decarboxylation of glutamate to γ-aminobutyric acid (GABA), which involves the take-up of a proton and the formation of one CO_2_ molecule (Castanie-Cornet et al. 1999). Via the glutamate/GABA antiporter (GadC), GABA is subsequently removed from the cell by exchanging it for glutamate (Castanie-Cornet et al. 1999; Cotter et al. 2001). In addition, all verrucomicrobial strains encode for the biosynthetic arginine decarboxylase (SpeA), which catalyzes the decarboxylation of arginine to agmatine, consuming a proton and producing one CO_2_ molecule (Richard and Foster 2004) (**Supplementary File S2**). The decarboxylated product can subsequently be removed from the cell by an antiporter (Slonczewski et al. 2009). Based on genome analyses only, it is not possible to decipher which genes encode the dedicated antiporters in verrucomicrobial methanotrophs. In *Escherichia coli*, GadB and GadC are inactive above pH 6.5, and both have pH optima of 4 to 5.5 (Ma et al. 2012; Pennacchietti et al. 2018). If the homologs in verrucomicrobial methanotrophs have similar pH optima, it is questionable whether this system is employed since even at very low external pH, the internal pH was shown not to decrease below pH 6 (Carere et al. 2021).

As opposed to neutrophilic microorganisms, acidophiles are constantly surrounded by an (extremely) acidic environment. If the intracellular proton concentration becomes too high, acidophiles could use cytoplasmic buffering and proton efflux pumps to restore the internal pH and the membrane potential (Mangold et al. 2013). Still, at a too low intracellular pH, proteins and DNA could become damaged due to a high concentration of protons (Baker-Austin and Dopson 2007). In verrucomicrobial methanotrophs, a large variety of chaperones are present that were shown to be produced in response to acid in neutrophiles, such as GroEL, GroES, DnaK and DnaJ (**Supplementary File S2**) (Carere et al. 2021; Frees et al. 2003; Len et al. 2004; Zanotti and Cendron 2010; Mols et al. 2010). These proteins are used by a wide range of organisms and assist in protein folding during times of stress, while damaged proteins are degraded by the enzyme Clp (Wawrzynow et al. 1996; Lemos and Burne 2002). In addition, RecA and UvrABCD were shown to be involved in acid-induced DNA repair (Hanna et al. 2001; Cappa et al. 2005; Jin et al. 2011; Das et al. 2015). Still, all these proteins are used in response to various stressors and therefore their presence does not necessarily indicate an adaptation to a low pH.

### *M. fumariolicum* SolV upregulates methane respiration genes in response to a low pH

Interestingly, none of the genes proposed to be involved in pH homeostasis in *M. fumariolicum* SolV are significantly upregulated when the bacterium is grown at pH 1.7 compared to growth at pH 3.0 (**Supplementary Table 1**). The absence of gene regulation when grown at these different pH values could have several explanations. It is possible that a shift in external pH from 3.0 to 1.7 (*i*.*e*., a 20-fold increase in proton concentration) is insufficient to modulate expression of the putative pH homeostasis genes. Indeed, the internal pH of *Methylacidiphilum* sp. RTK17.1 remains 6.5 when in an external pH of 1.5 to 3.0 and only fell below 6.5 at an external pH of 1 or lower (Carere et al. 2021). Feasibly, regulation of the genes proposed to be involved in pH homeostasis only occurs at even lower external pH values than those used here. Alternatively, it could be that the genes listed in **Supplementary Table 1** are not essential to *M. fumariolicum* SolV in terms of pH homeostasis. Although several genes were shown to be upregulated in response to a decrease in pH in acidophiles and neutrophiles, many proteins encoded by these genes could serve multiple purposes in the cell, which may explain their constitutive expression.

Remarkably, whereas genes predicted to be involved in pH homeostasis where not regulated at different pH values, several genes involved in respiration were heavily upregulated at pH 1.7 compared to pH 3.0 (**Table 2**; **Supplementary Table S1**; **Supplementary File S3**). *pmoBAC2*, one of three *pmo* operons in *M. fumariolicum* SolV encoding a particulate methane monooxygenase was upregulated 18-to 77-fold. This operon was shown to be highly upregulated under maximum growth conditions in batch cultures (Khadem et al. 2012). In addition, an *aa*_3_-type cytochrome *c* oxidase (Mfumv2_0388-92), an orphan subunit I of an *aa*_3_-type cytochrome *c* oxidase (Mfumv2_2181) and two genes involved in haem *a* synthesis (Mfumv2_1714-15) were upregulated (Svensson et al. 1993). *E. coli* was shown to upregulate proton-pumping respiratory complexes in response to acid stress (Slonczewski et al. 2009; Krulwich et al. 2011). Accordingly, the upregulation of genes involved in the aerobic oxidation of methane could be a response to an elevated ΔpH (*i*.*e*., an increase in the pH gradient component of the proton motive force due to a larger difference in proton concentration between the cytoplasm and the environment) and as such as mechanism to extrude excess protons (Siliakus et al. 2017; Carere et al. 2021). Alternatively, greater energetic investment are needed for cellular maintenance at low pH (Mangold et al. 2013). To further investigate these possibilities, detailed physiological studies on methane respiration at different pH values are desirable. Remarkably, the group 1d [NiFe] hydrogenase (Mfumv2_1562-66), known to oxidize H2 and feed electrons to the electron transport chain, is upregulated 5-fold in the absence of H2, suggesting it may work bidirectionally (Søndergaard et al. 1016; Mohammadi et al. 2017). In addition, three genes (Mfumv2_0526-7 and Mfumv2_0815) involved in the proton-consuming assimilatory conversion of sulfate (SO_4_^2-^) to sulfide (H_2_S) were upregulated.

**Table 2:** Regulation of *Methylacidiphilum fumariolicum* SolV genes of cells grown at pH 1.7 compared to cells grown at pH 3.0.

| ORF | Annotation | Regulation factor |
| --- | --- | --- |
| Mfumv2_0275 | Conserved exported protein of unknown function | 1.6 |
| Mfumv2_0388 | Cytochrome *c* oxidase polypeptide III | 2.2 |
| Mfumv2_0389 | Predicted small integral membrane protein | 1.9 |
| Mfumv2_0390 | Conserved protein of unknown function | 2.0 |
| Mfumv2_0391 | Conserved protein of unknown function | 1.7 |
| Mfumv2_0392 | Cytochrome *c* oxidase (*cyoA*) | 1.8 |
| Mfumv2_0393 | Conserved protein of unknown function | 1.6 |
| Mfumv2_0526 | Sulfate adenylyltransferase subunit 2 (*cysD*) | 1.8 |
| Mfumv2_0527 | Phosphoadenosine phosphosulfate reductase (cysH) | 1.9 |
| Mfumv2_0644 | Transposase | 2.1 |
| Mfumv2_0815 | Sulfite reductase [NADPH] hemoprotein beta-component (cysI) | 1.7 |
| Mfumv2_1257 | Conserved protein of unknown function | 1.6 |
| Mfumv2_1449 | Hemoglobin-like protein | 1.8 |
| Mfumv2_1462 | Pyrroloquinoline-quinone synthase (pqqC) | 1.7 |
| Mfumv2_1480 | Protein of unknown function | 2.1 |
| Mfumv2_1562 | Putative HupH hydrogenase expression protein | 4.2 |
| Mfumv2_1563 | Putative Ni/Fe-hydrogenase B-type cytochrome subunit (hupZ) | 5.1 |
| Mfumv2_1564 | Hydrogenase 1 large subunit (hyaB) | 5.3 |
| Mfumv2_1565 | Hydrogenase 1 small subunit (hyaA) | 5.0 |
| Mfumv2_1616 | Ribonuclease HIII | 1.6 |
| Mfumv2_1631 | Ribonuclease HIII (rnhC) | 1.6 |
| Mfumv2_1714 | Heme A synthase, cytochrome oxidase biogenesis protein (CtaA) | 2.5 |
| Mfumv2_1715 | Protoheme IX farnesyltransferase (ctaB) | 2.4 |
| Mfumv2_1758 | Multicopper oxidase family protein (cueO) | 1.6 |
| Mfumv2_1759 | Conserved protein of unknown function | 1.5 |
| Mfumv2_1794 | Methane monooxygenase subunit alpha (pmoB2) | 18.0 |
| Mfumv2_1795 | Methane monooxygenase subunit beta (pmoA2) | 29.3 |
| Mfumv2_1796 | Methane monooxygenase subunit gamma (pmoC2) | 76.8 |
| Mfumv2_1798 | Conserved protein of unknown function | 2.2 |
| Mfumv2_1799 | DNA-binding response regulator, NarL family REC-HTH domains | 2.2 |
| Mfumv2_1800 | ABC transport system, permease component YbhR | 1.7 |
| Mfumv2_1819 | Conserved protein of unknown function | 1.6 |
| Mfumv2_1832 | Globin domain | 2.2 |
| Mfumv2_2181 | Alternative cytochrome *c* oxidase subunit 1 (coxN) | 7.1 |
| Mfumv2_2188 | Conserved protein of unknown function | 2.9 |
| Mfumv2_2215 | Conserved membrane protein of unknown function | 1.6 |
| Mfumv2_2381 | Opacity protein or related surface antigen | 1.8 |
| Mfumv2_0170 | Outer membrane receptor protein, mostly Fe transport (cirA) | -26.7 |
| Mfumv2_0171 | Conserved protein of unknown function | -5.3 |
| Mfumv2_0553 | Iron-sulfur cluster insertion protein (ErpA) | -3.3 |
| Mfumv2_0556 | Pyridoxal phosphate-dependent enzyme | -1.8 |
| Mfumv2_1062 | Protein of unknown function | -1.5 |
| Mfumv2_1470 | Glutaredoxin-related protein | -1.7 |
| Mfumv2_2500 | Fe-S cluster scaffold complex subunit (SufC) | -1.7 |

Listed genes have a basemean higher than 4, an upregulation (positive values) or downregulation (negative values) factor higher or lower than 1.5 and an adjusted p-value $\leq 0.05$ . Numbers are averages of three technical replicates.

Several genes were downregulated at pH 1.7 compared to pH 3.0, among which the 27- and 5-fold downregulation of the outer membrane receptor protein (CirA) (Mfumv2_0170) and the adjacent gene with unknown function (Mfumv2_0171), respectively (**Table 2**). In *E. coli*, CirA was shown to be involved in the uptake of iron-chelating molecules (siderophores) (Hantke 1990). At lower pH, the solubility and bioavailability of metals generally increase, which may lead to downregulation of receptor protein expression to balance metal uptake. Interestingly, the outer membrane receptor protein CirA of *Methylacidiphilum fumariolicum* SolV shares sequence similarity, albeit it at 28% identity, with LutH, a protein thought to play a role in periplasmic lanthanide uptake (Groom et al. 2019; Roszczenko-Jasińska et al. 2020; Gorniak et al. 2023; Liu et al. 2025). Verrucomicrobial methanotrophs, including *M. fumariolicum* SolV, rely on lanthanides as essential cofactors for the activity of the methanol dehydrogenase XoxF (Pol et al. 2014; Schmitz et al. 2021b). While considerable progress has been made in identifying lanthanide-binding and -uptake systems in proteobacterial methylotrophs, the molecular mechanisms underlying lanthanide acquisition in verrucomicrobial methanotrophs remain poorly understood. Further experimental investigation is required to identify and characterize novel proteins involved in lanthanide binding and uptake in these extremophiles.

## Conclusion

In conclusion, in this study we show the fatty acid compositions of three strains of mesophilic *Methylacidimicrobium* strains for the first time. In general, diverse membrane fatty acid compositions were observed for verrucomicrobial methanotrophs, suggesting that multiple adaptive strategies exist to modulate membrane properties for survival under low pH and high-temperature conditions. The membranes of mesophilic *Methylacidimicrobium* strains consist on average for more than a quarter of non-saturated fatty acids, which are virtually absent in thermophilic *Methylacidiphilum* strains. The level of unsaturation could be explained by mechanisms to preserve appropriate membrane fluidity at different growth temperatures. Indeed, the thermophile *Methylacidiphilum fumariolicum* SolV was shown to synthesize monounsaturated C18 fatty acids in response to a decrease in temperature of 10 °C. Still, the membrane fatty acid composition of *Methylacidiphilum fumariolicum* SolV and *Methylacidimicrobium cyclopophantes* 3B grown at almost identical pH and temperature are not in the least identical. Hence, even for closely related strains, multiple ways exist to thrive at elevated temperatures at extremely low pH. The fatty acid biosynthesis pathway in verrucomicrobial methanotrophs is conserved, although *Methylacidimicrobium* strains may possess a more versatile pathway to synthesize unsaturated fatty acids. Many genes used by neutrophiles for pH homeostasis were detected in verrucomicrobial methanotrophs and expressed in *M. fumariolicum* SolV. However, a decrease in pH from 3.0 to 1.7 did not result in regulation of these genes. *M. fumariolicum* SolV primarily responded to a decrease in pH by the upregulation of genes involved in methane respiration pathway, presumably to pump out excess protons. This study highlights the importance of validated predictions through comparative genomics with experimental studies.

## Declarations

### Ethics statement

This article does not contain any studies with human participants or animals performed by any of the authors.

### Data availability

Data is provided within the manuscript or supplementary information files.

### Supplementary Material

The Supplementary Material for this article can be found online.

