## Supplementary Information for "Adaptation and response of verrucomicrobial methanotrophs to heat and acidity"

**Supplementary Table S1**: Expression of genes predicted to be involved in pH homeostasis in *Methylacidiphilum fumariolicum* SolV when grown at pH 3.0 and pH 1.7. TPM: Transcripts Per Million. Standard deviations are indicated ($n$ = 3). ORF: open reading frame.

| ORF | Annotation | TPM (pH 3.0) | TPM (pH 1.7) |
|---|---|---|---|
| ***Potassium influx*** | | | |
| Mfumv2_0028 | Potassium-transporting ATPase subunit KdpC | 41 ± 10 | 35 ± 5 |
| Mfumv2_0029 | Potassium-transporting ATPase subunit KdpB | 58 ± 6 | 50 ± 2 |
| Mfumv2_0030 | Potassium-transporting ATPase subunit KdpA | 46 ± 4 | 44 ± 5 |
| Mfumv2_1639 | Potassium channel protein Kch | 79 ± 10 | 57 ± 4 |
| Mfumv2_1842 | Regulator of kdp operon KdpE | 97 ± 10 | 87 ± 8 |
| Mfumv2_1843 | Sensor histidine kinase KdpD | 113 ± 15 | 105 ± 6 |
| ***Proton efflux*** | | | |
| Mfumv2_0748 | Cation:proton antiporter KefB | 23 ± 6 | 29 ± 4 |
| Mfumv2_1662 | Cation:proton antiporter KefB | 45 ± 5 | 41 ± 4 |
| Mfumv2_2336 | Sodium-translocating pyrophosphatase HppA | 238 ± 37 | 177 ± 14 |
| ***Proton consuming mechanisms through decarboxylation*** | | | |
| Mfumv2_0149 | Glutamate decarboxylase GadA/GadB | 208 ± 36 | 206 ± 15 |
| Mfumv2_0264 | Biosynthetic arginine decarboxylase SpeA | 303 ± 66 | 277 ± 33 |
| ***Protein and DNA protection and repair*** | | | |
| Mfumv2_0146 | ATP-dependent chaperone ClpB | 587 ± 46 | 538 ±16 |
| Mfumv2_0624 | Excinuclease ABC subunit UvrB | 264 ± 21 | 245 ± 12 |
| Mfumv2_1105 | ATP-dependent helicase UvrD | 77 ± 10 | 69 ± 4 |
| Mfumv2_1808 | Excinuclease ABC subunit UvrC | 90 ± 9 | 106 ± 7 |
| Mfumv2_1913 | Recombinase RecA | 532 ± 34 | 544 ± 17 |
| Mfumv2_2124 | Excinuclease ABC subunit UvrA | 87 ± 3 | 82 ± 4 |
| Mfumv2_2206 | ATP-dependent Clp protease ATP-binding subunit | 1572 ± 74 | 1529 ± 73 |
| Mfumv2_2307 | Chaperonin GroEL | 6159 ± 474 | 5162 ± 158 |
| Mfumv2_2308 | Co-chaperone GroES | 4813 ± 139 | 4038 ± 65 |
| Mfumv2_2309 | Molecular chaperone DnaK | 2715 ± 138 | 2349 ± 172 |
| Mfumv2_2371 | Molecular chaperone DnaJ | 929 ± 102 | 865 ± 11 |

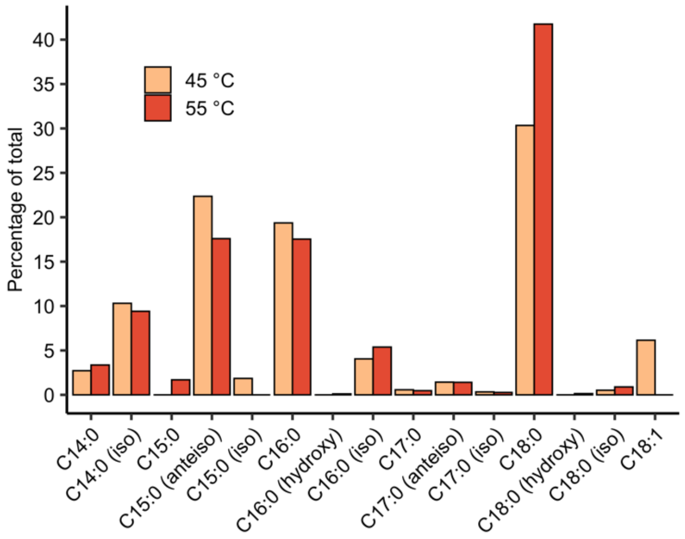


**Supplementary Figure 1**: Fatty acids of *Methylacidiphilum fumariolicum* SolV cells grown during steady state in chemostats at pH 3.0 at 45 °C and 55 °C. anteiso: *anteiso*-fatty acids; hydroxy: hydroxyl fatty acids; iso: *iso*-fatty acids.
